# *In situ* architecture of the ciliary base reveals the stepwise assembly of IFT trains

**DOI:** 10.1101/2021.10.17.464685

**Authors:** Hugo van den Hoek, Nikolai Klena, Mareike A. Jordan, Gonzalo Alvarez Viar, Miroslava Schaffer, Philipp S. Erdmann, William Wan, Jürgen M. Plitzko, Wolfgang Baumeister, Gaia Pigino, Virginie Hamel, Paul Guichard, Benjamin D. Engel

## Abstract

The cilium is an antenna-like organelle that performs numerous cellular functions, including motility, sensing, and signaling. The base of the cilium contains a selective barrier that regulates the entry of large intraflagellar transport (IFT) trains, which carry cargo proteins required for ciliary assembly and maintenance. However, the native architecture of the ciliary base and the process of IFT train assembly remain unresolved. Here, we use *in situ* cryo-electron tomography to reveal native structures of the transition zone region and assembling IFT trains at the ciliary base. We combine this direct cellular visualization with ultrastructure expansion microscopy to describe the front-to-back stepwise assembly of IFT trains: IFT-B forms the backbone, onto which IFT-A, then dynein-1b, and finally kinesin-2 sequentially bind before entry into the cilium.

**One Sentence Summary:** Native molecular structure of the ciliary transition zone and hierarchical order of IFT assembly visualized within *Chlamydomonas* cells.

## Main Text

Cilia (also called flagella) are evolutionarily-conserved eukaryotic organelles that extend from the cell surface and carry out a wide variety of functions, including cell motility, fluid flow generation, sensing, and signaling (*1, 2*). Because cilia perform many important roles, defects in cilia structure and function are linked to pleiotropic human diseases, called “ciliopathies” (*3*). The cilium consists of an axoneme of microtubule doublets (MTDs), which extend from the microtubule triplets (MTTs) of the centriole (also known as basal body) and are sheathed in a ciliary membrane. At the base of the cilium, a specialized region known as the transition zone (TZ) gates entry and exit of both membrane-bound and soluble ciliary proteins (*4–6*). Defects in TZ proteins lead to altered cilium composition and cause diseases including Nephronophthisis (NPHP) and Meckel-Gruber syndrome (MKS) (*7, 8*). However, a molecular structure of the TZ has remained elusive.

Among its many gating functions, the TZ is thought to regulate entry of intraflagellar transport (IFT), the kinesin- and dynein-driven bidirectional traffic of axonemal cargos and membrane proteins between the ciliary base and tip (*9, 10*). IFT is required for assembly and maintenance of the cilium, as well as for mediating many of its signaling functions (*11*). Fluorescence recovery after photobleaching (FRAP) experiments have revealed that a pool of IFT and motor proteins resides at the ciliary base for several seconds before entry into the cilium (*12, 13*). Furthermore, different proteins in the pool recover fluorescence at distinct rates, suggesting ordered recruitment. However, it remains to be characterized how these IFT proteins and motors assemble into the elaborate multi-megadalton trains observed running along the ciliary axoneme (*14*). Here, we combine *in situ* cryo-electron tomography (cryo-ET) (*15*) with ultrastructure expansion microscopy (U-ExM) (*16*) of the green alga *Chlamydomonas reinhardtii*, a classic model for cilia research, to reveal native TZ structures and stepwise assembly of IFT trains at the ciliary base.

### Native structure of the *Chlamydomonas* transition zone

Vitrified *Chlamydomonas* cells were thinned with a focused ion beam (*17*) then imaged by cryo-ET, yielding 19 tomograms of the ciliary base inside the native cellular environment. Focusing on the TZ, we observed several different structures attached to the MTDs (Fig. 1A-B), which we resolved in molecular detail by subtomogram averaging (Figs. 1C-H, S1). The proximal ~180 nm of the TZ is occupied by peripheral Y-links (Fig. 1, turquoise) and luminal stellate fibers (Fig. 1, purple), which resemble a 9-pointed star in cross-section (*18*). Our structure reveals that the stellates form a helical cylinder with a 6-start helix and a pitch of 49.2 nm (Fig. 1E and H, S2). Interestingly, this matches the pitch of the inner scaffold, a 3-start helical cylinder in the lumen of the centriole (*19*). The nine points of the stellate star bind MTDs at protofilament A3, with a longitudinal periodicity of ~8.1 nm along each MTD, which matches the 8.2 nm length of one tubulin dimer (Figs. 1F, S2). TZ stellates have been observed in certain algae and early-evolving land plants, and have a contractile function that is believed to trigger ciliary abscission (*20*).

**Figure 1.**
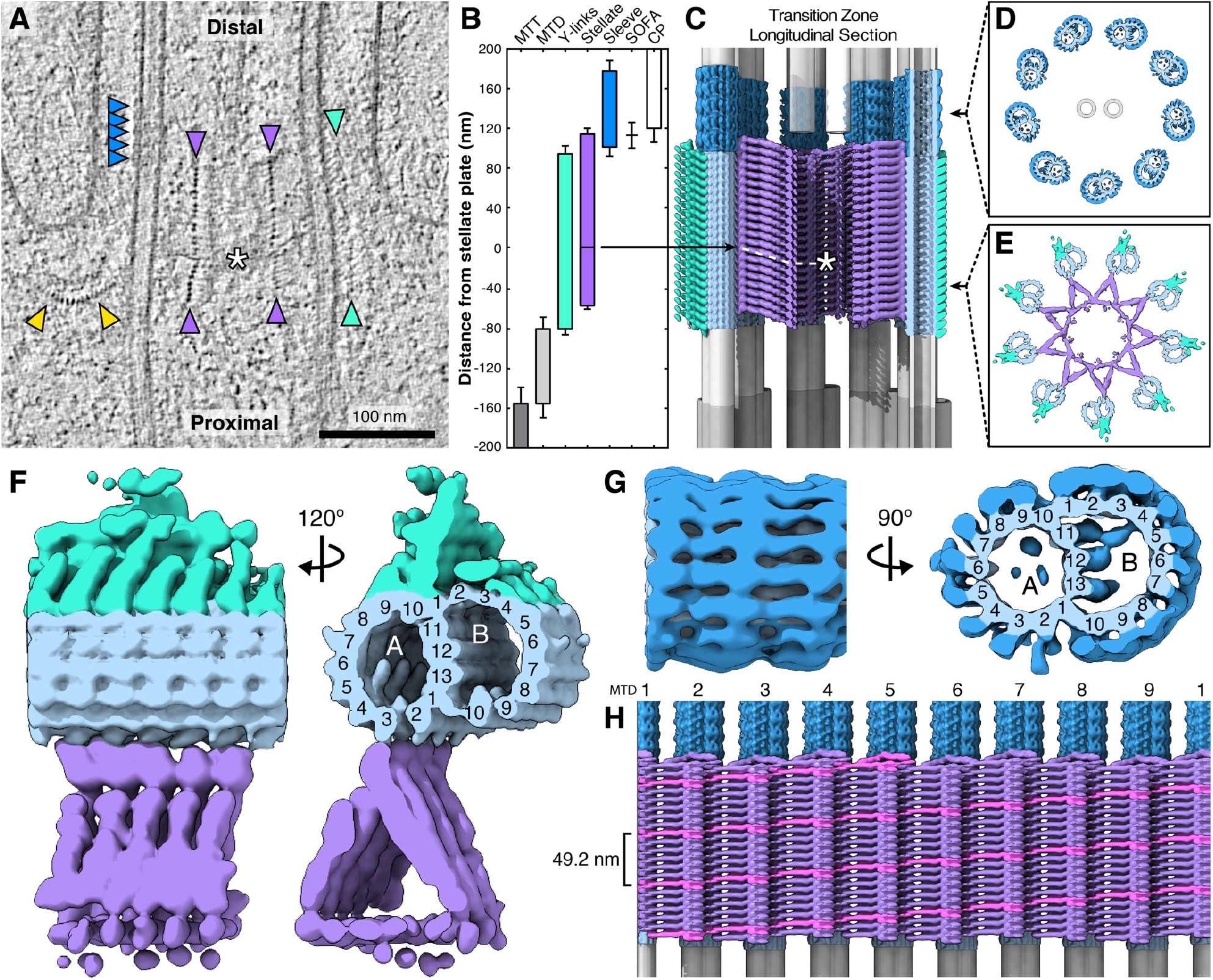
Cryo-ET structure of the ciliary transition zone inside *Chlamydomonas* cells. **(A)** 2D slice through a cryo-electron tomogram, showing a TZ in longitudinal cross-section. Purple: stellate fibers, turquoise: Y-links, dark blue: MTD helical sleeve, white asterisk: stellate plate, yellow: IFT (see Fig. 2). Scale bar: 100 nm. **(B-G)** Composite model of the TZ, combining averages of the stellate (purple), Y-links (turquoise), MTD sleeve (dark blue), and associated MTD (light blue), with schematic renderings of MTTs (dark grey), MTDs (light grey), and the central pair (CP, white). **(B)** Position along the TZ occupied by different structures. Distances are measured relative to the stellate plate (“0 nm” origin point marked with arrow). SOFA: “site of flagellar autotomy”, where the cilium is cleaved (see Fig. S3). Error bars: standard deviation. **(C)** Longitudinal section view of the complete composite model, assembled according to the measured lengths and positions of each component, with 21 Y-link repeats, 21 stellate repeats (7 proximal of the plate, 14 distal), and 5 MTD sleeve repeats. The model shows straight MTTs and MTDs, but as seen in panel A and quantified in (*19*), the centriole is actually a slightly convex barrel. **(D-E)** Cross-section views through the indicated regions of the composite model, showing **(D)** MTDs encased in the helical sleeve, with the CP in the middle, and **(E)** the nine-pointed stellate cylinder attached to MTDs decorated with Y-links. (**F-G)** Side and cross-section views a single MTD attached to **(F)** stellate fibers and Y-links, and **(G)** the helical sleeve. Protofilaments of the A- and B-microtubules are numbered. **(H)** Unrolled composite model, viewed from the center of the TZ, looking outward toward the inner wall of the stellate cylinder. One continuous helical density of the stellate’s six-start stellate helix is marked in pink (pitch: 49.2 nm). MTDs are numbered.

Y-links are present in many species. They connect MTDs to the ciliary membrane and contain NPHP- and MKS-family proteins that are proposed to gate transport into and out of the cilium (*7, 8*). Our structure reveals a broad density attaching the Y-links to the MTD, spanning protofilaments A9-10 and B1-4, with a longitudinal repeat of ~8.3 nm (Fig. 1F, S2). However, the outer densities of the Y-links that connect to the ciliary membrane were not resolved, likely due to flexibility of these filamentous structures. Interestingly, we observed that the Y-links extend along the MTDs proximal of the region where the ciliary membrane aligns with the axoneme (Fig. 1A). Thus, connection to the membrane is not a prerequisite for MTD Y-link decoration, suggesting that the Y-links may have more functions in addition to membrane-axoneme tethering.

Immediately distal to the stellates and Y-links, we found a distinct helical density completely decorating the MTDs (Fig. 1, dark blue), which to our knowledge has not been described before. This helical “sleeve” spans 76 ±6 nm along the MTDs, with a periodicity of ~16.4 nm (Fig. 1G, S2). To gain hints into the function of this structure, we also analyzed tomograms of basal bodies isolated immediately following ciliary shedding (Fig. S3B). The *in situ* position of the MTD sleeve overlaps with the ciliary cleavage site observed in isolated basal bodies, also known as the site of flagellar autotomy (SOFA) (*21*) (Fig. 1B). Consistent with this finding, the MTD sleeve was absent from the distal ends of these basal bodies. The sleeve was also not observed on the proximal ends of isolated cilia (*22*). We therefore hypothesize that this structure might help regulate axoneme severing and is lost during the process. Since the sleeve density covers every MTD protofilament, it should sterically hinder the attachment of molecular motors, and thus, may also play a role in regulating IFT entry or exit.

### Anterograde IFT trains assemble in the cytoplasm prior to ciliary entry

It has been known for decades that a pool of IFT proteins is localized near the base of the cilium (*23, 24*), but the structural organization of this basal pool has remained a mystery. In our tomograms, we observed filamentous strings of particles with one end attached to the TZ and the other end splayed into the cytosol (Fig. 2A). These strings appeared to consist of three layers, each with a different shape and periodicity (Fig. 2B-C). Iterative 3D alignment of particles picked along each layer yielded subtomogram averages that we combined to produce a composite molecular structure of the strings (Figs. 2D, S4). Comparison to a previously published cryo-ET structure of mature anterograde IFT trains within the cilium (*14*) confirmed that the cytosolic strings are indeed IFT trains. Our structure of cytosolic trains was similar to the anterograde train structure, enabling us to assign densities to IFT-B complexes, IFT-A complexes, and dynein-1b motors, which have longitudinal periodicities of ~6 nm, ~11 nm, and ~18 nm, respectively (Fig. S5). As was shown for anterograde IFT, we observed that dynein-1b is loaded onto cytosolic IFT trains as a cargo in an autoinhibited state (*14, 25*). We therefore conclude that much of the IFT pool at the ciliary base consists of TZ-tethered anterograde IFT trains that are undergoing assembly prior to entry into the cilium. We did not notice obvious retrograde trains at the ciliary base, implying that they rapidly disassemble, perhaps even before exiting the TZ.

**Figure 2.**
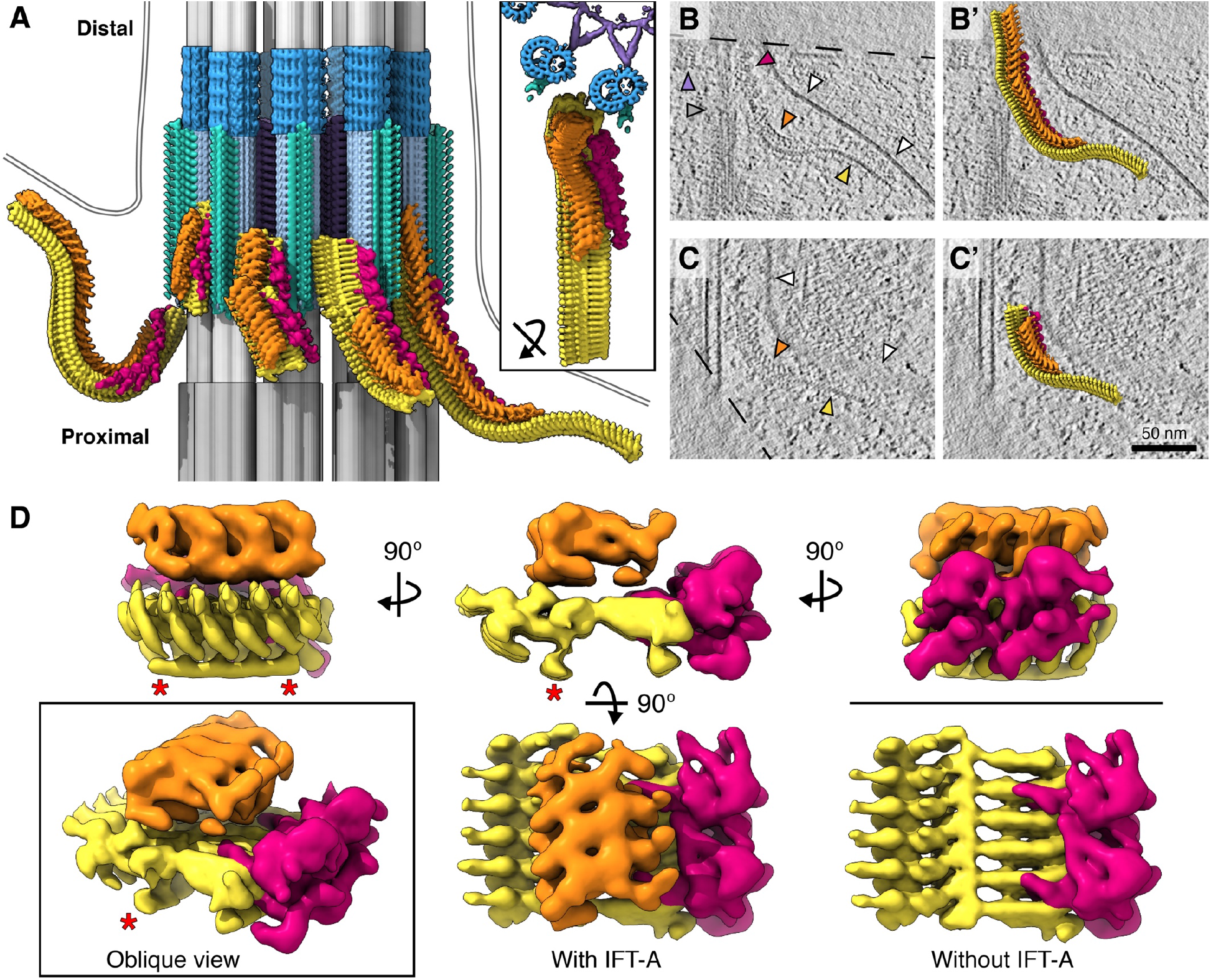
Cryo-ET structures of assembling IFT trains attached to the transition zone. **(A)** Composite structures of assembling IFT trains (yellow: IFT-B; orange: IFT-A; red: dynein-1b) engaging the ciliary TZ (colored as in Fig. 1). A schematic membrane has been added (grey double lines). Subtomogram averages have been mapped back into their positions inside a representative tomogram. **Inset:** rotated top view of rightmost IFT train from the main panel. **(B-C)** 2D slices through tomograms, showing assembling IFT trains, and **(B’-C’)** overlaid 3D train structures, with their front ends attached to the TZ. In panel B, note the double AAA+ ATPase rings of the dynein-1b motors. Arrowhead colors correspond to panel A, and additional white arrowheads indicate the membrane. Scale bar: 50 nm. **(D)** Different views on the cytosolic IFT train composite subtomogram average. Colored as in panel A. Red asterisks indicate the extra IFT-B density, which was not observed on mature trains inside cilia (see Fig. S6).

We observed two notable differences between assembling and mature IFT trains. First, assembling trains are flexible and display regions of high curvature (Figs. 2A-C, S5), whereas trains in the cilium have an extended straight conformation (Fig. S5), which is likely maintained by interactions with cargos and the ciliary membrane. Second, assembling IFT-B has a prominent extra density on the side opposite IFT-A (Figs. 2D, S6; red asterisks), close to IFT-B’s proposed kinesin-2 binding site (*14*). In regions where IFT trains are bound to the TZ, this extra density is positioned immediately adjacent to the MTD. The density is not part of the MTD structure itself, since it is also present on IFT-B in the sections of trains that dangle into the cytosol (Fig. S6E). One possibility is that this density acts a molecular brake that prevents entry of IFT trains into the cilium until they are fully assembled and loaded with cargo. However, in a few cases, we also observed the density on trains along the axoneme just distal of the TZ (Fig. S6D). Understanding the interactions and function of the extra IFT-B density will require a high-resolution IFT structure that reveals the position of each IFT protein within the train.

### IFT trains assemble front-to-back in a stepwise fashion

*In situ* cryo-ET combines native molecular structures with precise cellular localization, enabling us to plot the spatial relationship between assembling IFT trains and the TZ (Fig. 3A-B). Although many IFT trains in our dataset were not completely contained within the thin FIB-milled cellular sections, 40 trains were largely intact and could be analyzed. The front segments of IFT trains attach to MTDs (but never to the MTTs of the centriole) and then continue in the distal direction along these microtubule tracks, while the back regions curve away from the TZ into the cytosol. The front ends of IFT trains were frequently observed at the Y-links, but only very rarely further distal at the MTD sleeve (Fig. 3A-B). By mapping the IFT-B, IFT-A, and dynein-1b structures along the IFT trains (as shown in Fig. 2A), we visualized the sequential order of train assembly. Anterograde trains always contain IFT-B, which forms a backbone scaffold upon which other components are attached. Thus, we plotted the train assembly state relative to each IFT-B subunit. While we observed variability between individual trains (Fig. 3A), the cumulative plot shows a clear distribution of IFT train components (Fig. 3B). The front parts of the trains are more complete, whereas the back segments are in a less assembled state, with IFT-B extending the furthest, followed by IFT-A and then dynein-1b. No gaps were observed in the regions occupied by IFT-A and dynein-1b, suggesting that these components linearly oligomerize upon the IFT-B scaffold. These results support a front-to-back mode of assembly, where IFT-B forms a backbone onto which IFT-A and then dynein-1b are sequentially added.

**Figure 3.**
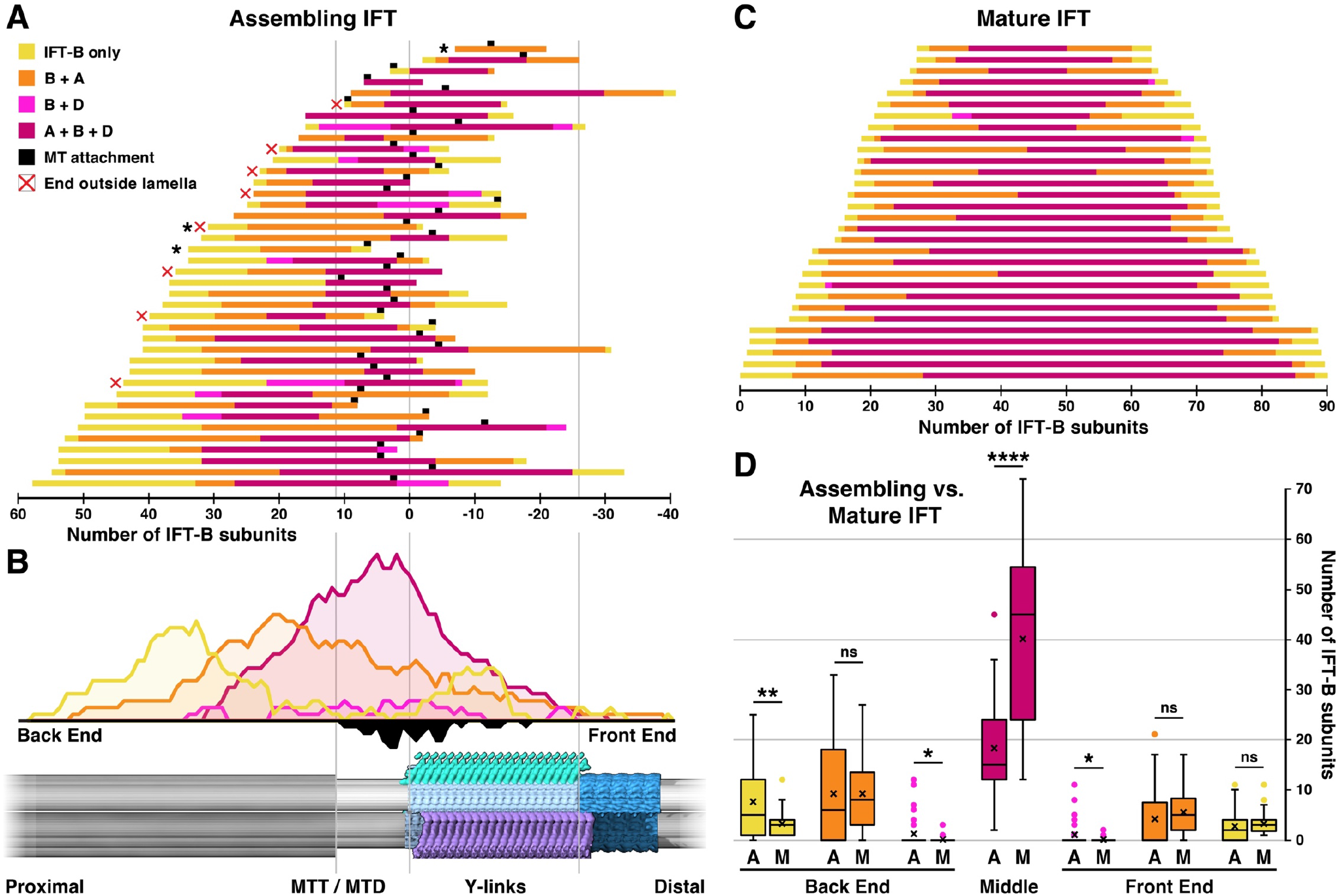
IFT trains undergo stepwise assembly from front to back. **(A-B)** Assembly states of IFT trains relative to structures of the TZ (shown at the bottom of panel B). Distances are measured in number of IFT-B subunits relative to the onset of the Y-links (defined as the “0 IFT-B” origin point). **(A)** Position and train assembly state, measured for each IFT-B subunit of each assembling IFT train. Assembly states (color code in legend) were determined based on occupancy of IFT-A and/or dynein-1b on the IFT-B backbone. Black squares indicate train attachment points, i.e. the most proximal IFT-B subunit that is close enough to the MTD (<20 nm) to be considered bound to the microtubule. The three trains marked with black asterisks lack a “middle” (no dynein is bound), and thus a “back end” could not be distinguished from a “front end”, so they were omitted from analysis in panel D. Of the 70 trains in our dataset, 30 had front ends cropped by the FIB milling, and therefore were not quantified, as their starting point relative to the TZ could not be determined. Eight more trains had cropped back ends (indicated with red “x”), but were included in the analysis. **(B)** Cumulative plot of all assembling IFT trains in panel A, with positions in relation to the TZ (shown at bottom). Colored curves above the line show the summed abundance of each IFT assembly state. The black curve below the line shows the distribution of MTD attachment points, summed from the black squares in panel A. **(C)** Distribution of IFT train lengths and completeness of assembly (colored as in panel A) for mature anterograde trains found in the cilium (*14*). **(D)** Comparison of the abundance of assembly states at the front and back ends of assembling (“A”) vs. mature (“M”) trains (i.e., trains in panel A vs. trains in panel C). Box: median and 25%–75% percentiles; X: mean; whiskers: 1.5× interquartile range; points: outliers. Statistical significance assessed by unpaired t-test (****, P < 0.0001; **, P = 0.002; *, P = 0.03; ns, not significant, P > 0.05). Because the backs of some assembling trains in this analysis were clipped by the FIB milling (“x” in panel A), the “IFT-B only” region of assembling trains should be slightly more pronounced than the quantification shown here.

Next, we compared the assembling cytosolic trains to mature anterograde trains within the cilium (Fig. 3C-D). Mature trains exhibited a continuum of lengths from 223 to 558 nm, comprising 36 to 90 IFT-B subunits with an average of 62 ±16 IFT-B. In comparison, the longest assembling train we observed contained 88 IFT-B subunits, and the average length was 44 ±17 IFT-B. Therefore, we conclude that just one IFT train is assembled at a time per MTD. Cytosolic and ciliary IFT trains have similarly incomplete front ends, whereas the incomplete segment at the back of cytosolic trains is longer, as these regions are still undergoing assembly (Fig. 3D). Perhaps as IFT trains enter the cilium, part of the incompletely assembled back end is left behind at the TZ to nucleate the front of the next train. As assembly of IFT-A and dynein-1b proceed front-to-back, the front end of the new train would remain incomplete, as we see in mature anterograde trains inside the cilium (Fig. 3C).

### Kinesin is loaded onto IFT trains close to the transition zone

The kinesin-2 anterograde IFT motor is difficult to identify by cryo-ET due to its relatively small size and flexible structure. Therefore, we turned to U-ExM super-resolution microscopy to examine the occupancy of kinesin-2 on assembling trains. We imaged three strains containing fluorescently-tagged IFT proteins: a kinesin-2 component (KAP-GFP), a dynein-1b component (d1bLIC-GFP), and an IFT-B component (IFT46-YFP). All three strains revealed train-like filamentous strings at the ciliary base, with their distal ends in close proximity to the TZ (marked by its lack of polyglutamylated tubulin, Fig. S7) and their proximal ends curving into the cytoplasm (Fig. 4A-E). Using semi-automated tracing, we plotted the 3D trajectories of each IFT string relative to the TZ (Fig. 4F-G). IFT46 traces were longer than d1bLIC traces, consistent with our cryo-ET observations. KAP traces were the shortest and were localized close to the TZ. Thus, our combined cryo-ET and U-ExM measurements support a consensus model of IFT train assembly: IFT-B first forms the backbone, which scaffolds the sequential attachment of IFT-A, dynein-1b, and finally, kinesin-2 close to the TZ (Fig. 4H).

**Figure 4.**
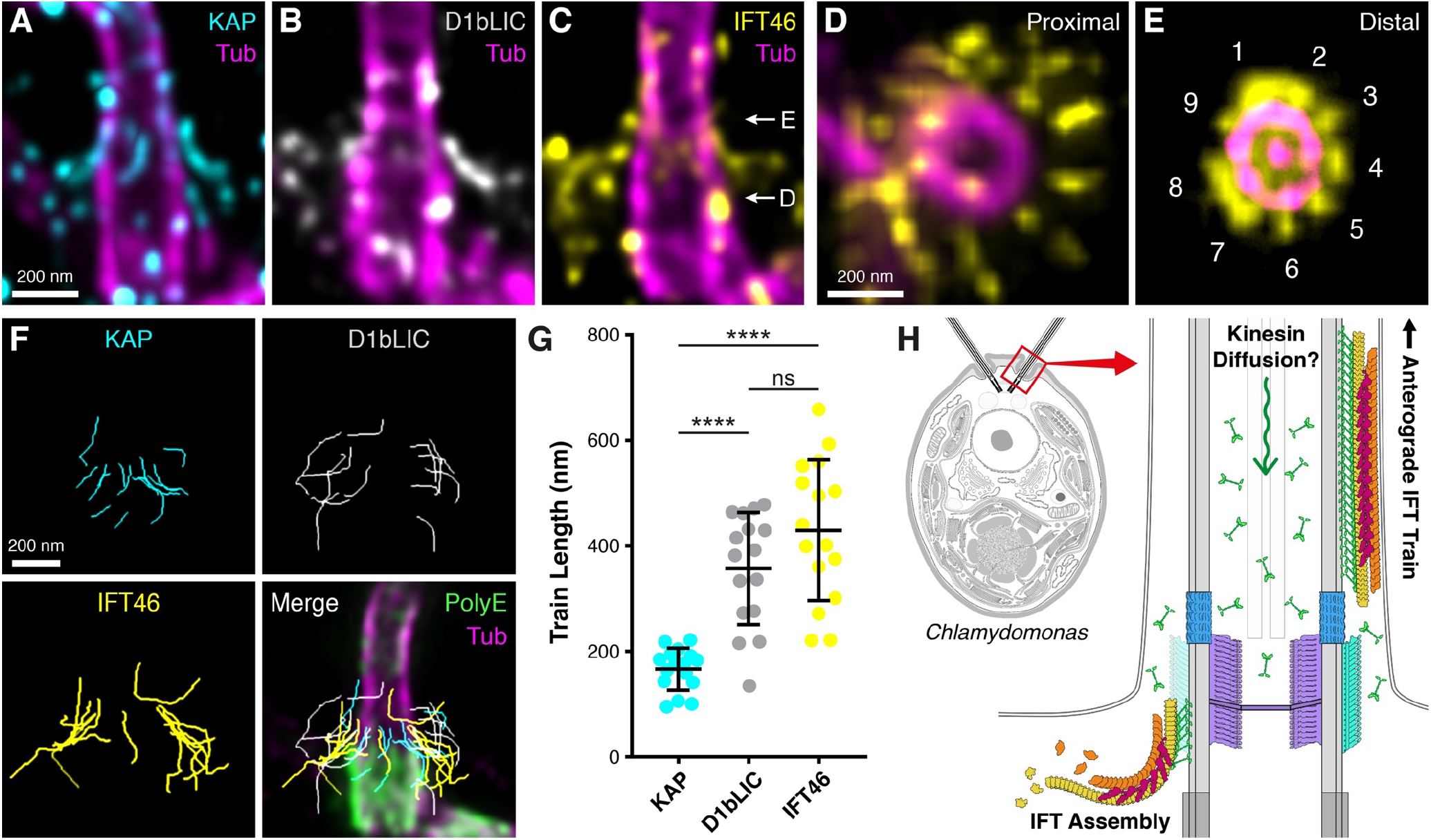
Assembly of anterograde IFT trains examined by U-ExM. **(A-C)** Longitudinal views of assembling IFT trains at the ciliary base, visualized by antibody staining and U-ExM. **(A)** KAP-GFP (cyan). **(B)** D1bLIC-GFP (gray). **(C)** IFT46-YFP (yellow). Tubulin staining is magenta. White arrows indicate the approximate region of top view z-projections presented in panels D and E. **(D, E)** Top view z-projections displaying 9-fold occupancy of IFT trains at the proximal TZ (D) and more distal TZ (E), confirmed by the presence of the central pair. **(F)** 3D traces of individual IFT trains stained for KAP, D1bLIC, and IFT46. Bottom right panel, all traces merged and overlaid on single ciliary base stained for polyglutamylated tubulin (PolyE, green) and tubulin (magenta). The gap in PolyE staining marks the TZ (see Fig. S7). **(G)** Measured lengths of 3D traces. N= 16 trains per stain, from 2 independent experiments. Statistical significance assessed by unpaired t-test (****, P < 0.0001; ns, not significant, P > 0.05). **(H)** Consensus model of IFT train assembly from cryo-ET and U-ExM measurements. New IFT-B subunits are added to the back of the train, forming a scaffold that is sequentially loaded with IFT-A, dynein-1b, and finally kinesin-2 close to the TZ. U-ExM of KAP-GFP indicates that kinesin-2 may diffuse from the ciliary tip to base through the axoneme lumen (see Fig. S8). Scale bars: 200 nm.

U-ExM revealed an apparent 9-fold occupancy of IFT around the TZ (Fig. 4D-E), consistent with recent observations in *Tetrahymena* (*26*). Given that each of these 9 signals corresponds to a single assembling IFT train (Fig. 3), and that trains enter the cilium with a frequency of ~1 train/sec (*27, 28*), we predict that trains should linger at the ciliary base for an average of 9 seconds during assembly. Consistent with this hypothesis, FRAP analysis of the *Chlamydomonas* basal pool showed that several GFP-tagged IFT proteins require ~9 seconds to recover peak fluorescence (*12*). Regardless, ciliary entry of IFT trains appears to be stochastic rather than sequential, with an avalanche-like relationship between lag time and train size (*29*).

The big question remains: what regulates entry of IFT into the cilium? In other words, what prevents the front end of the train from entering the cilium as assembly proceeds toward the incomplete back of the train? Maybe the extra IFT-B density we observed acts as a molecular brake, or perhaps IFT entry is dictated by kinesin-2 loading at the TZ. Interestingly, we observed a strong KAP-GFP signal inside the axoneme but excluded from the TZ and centriole (Fig. S8). One possible explanation could be that kinesin-2 strongly binds central pair microtubules, as an antibody raised against a *Drosophila* kinesin motor domain was shown to decorate central pairs isolated from *Chlamydomonas* (*30*). However, this hard to envision because this pair of microtubules is almost completely ensheathed in central apparatus proteins (*31*). Alternatively, the KAP-GFP localization may suggest that kinesin-2 diffuses back to the ciliary base through the axoneme lumen. The diffusion of KAP-GFP from the ciliary tip has been well documented in *Chlamydomonas* (*27, 32, 33*). In addition, high-speed super-resolution fluorescence microscopy showed that KIF17, the mammalian homolog of kinesin-2, diffuses within the axoneme lumen of primary cilia (*34*). It has been proposed that diffusive return of kinesin-2 from the ciliary tip could serve as a “ruler” that regulates the entry of new IFT trains (*33, 35*). However, FRAP experiments indicate that the cilium is an open system for kinesin-2; new trains bind fresh kinesin from the cytosol in addition to recycled kinesin from the cilium (*12*). Regardless, it is clear from our study that kinesin-2 loading is limited to the front regions of assembling trains (Fig. 4). A simple explanation could be that assembly factors required to load kinesin-2 onto trains are localized to the TZ. However, the architecture of the TZ itself may also play a role. Perhaps the stellates block proximal diffusion of kinesin-2, redirecting it from the axoneme lumen to load onto the front ends of assembling IFT trains (Fig. 4H). Indeed, IFT-bound kinesin-2 likely mediates train attachment to MTDs at the TZ (Fig. 3A-B, black). When sufficient kinesin-2 is bound to the front of the train, these motors may begin to drag the train in the distal direction through the dense matrix surrounding the Y-links that form the cilium’s diffusion barrier. As the train moves forward, additional kinesin-2 would be able to bind further back along the train as these parts enter the TZ. Eventually enough kinesin-2 would be bound along the train to propel it past the TZ structures and onward to start its anterograde journey through the cilium.

## Supplementary Figures

**Figure S1.**
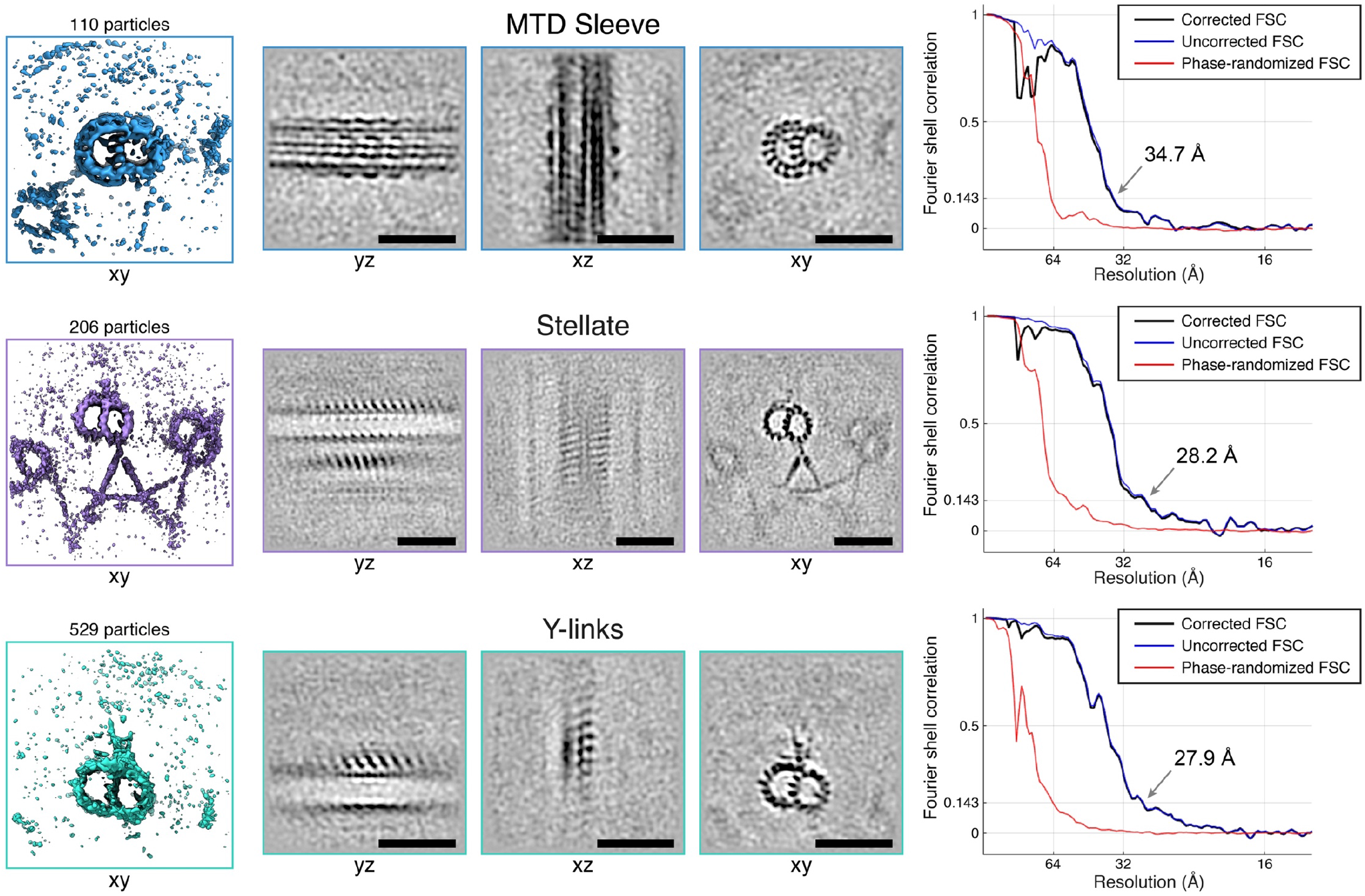
Individual subtomogram averages focused on different TZ structures. Subtomogram averages focused on individual components of the transition zone: the MTD helical sleeve (top row), stellate (middle row), and Y-links (bottom row). These averages were fused to generate the TZ model shown in Fig. 1. Left panels: 3D isosurface views, with the number of subvolumes (particles) in each average indicated. Central panels: 2D greyscale slices through the averages from three orientations (scale bars: 50 nm). Right panels: Fourier shell correlation curves showing the determined resolution of each average.

**Figure S2.**
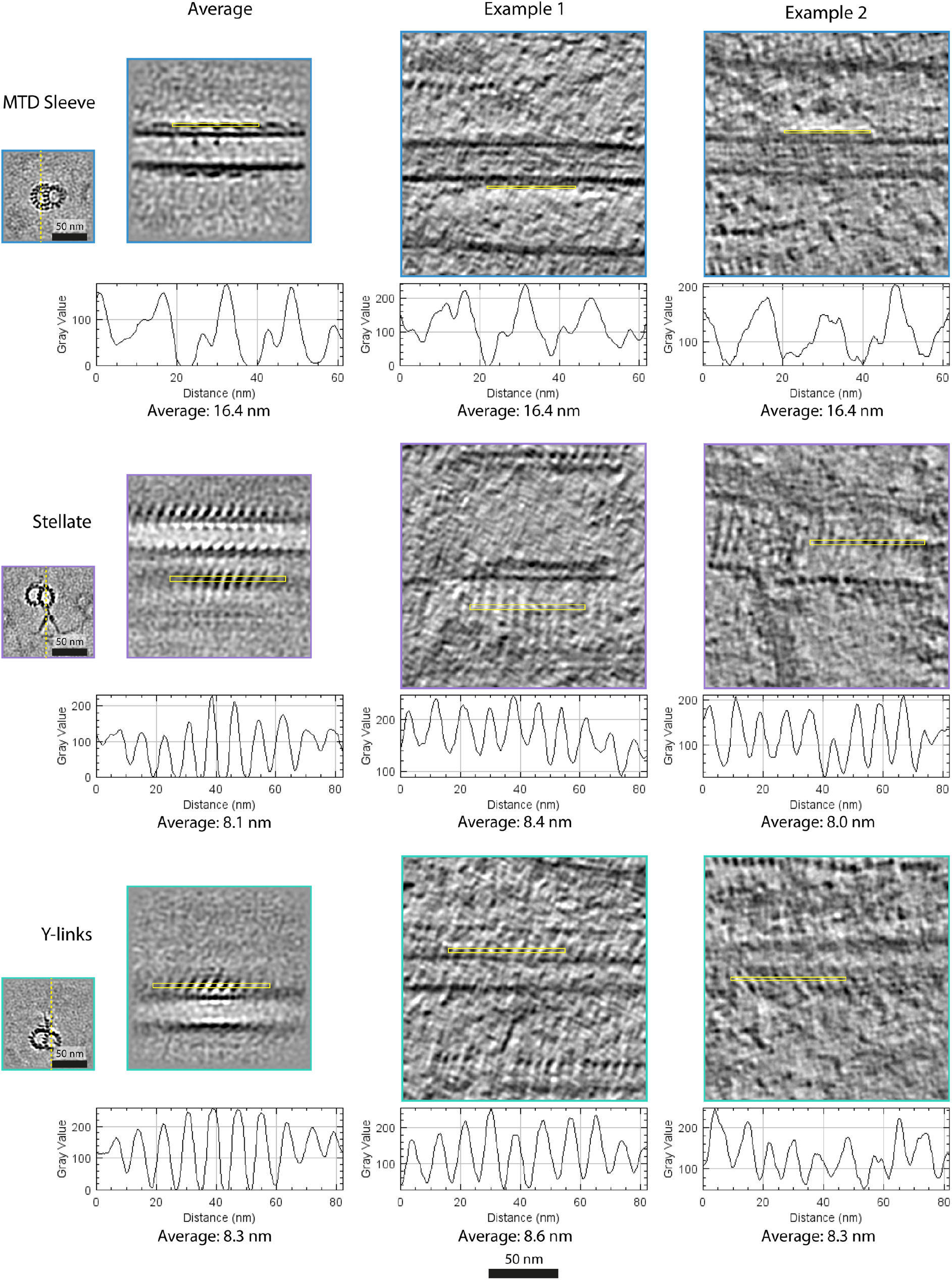
Periodicity of each TZ structure measured from greyscale 2D slices. For each structure, subtomogram averages are shown on the left, and two examples of raw tomograms are shown on the right. The orientations of the longitudinal slices are indicated with dashed yellow lines in the cross-section views on the far left. Line scans along the yellow boxes in the longitudinal views were generated with FIJI software (*36*). Corresponding intensity plots are shown below each longitudinal slice, along with the average periodicity measured from each plot.

**Figure S3.**
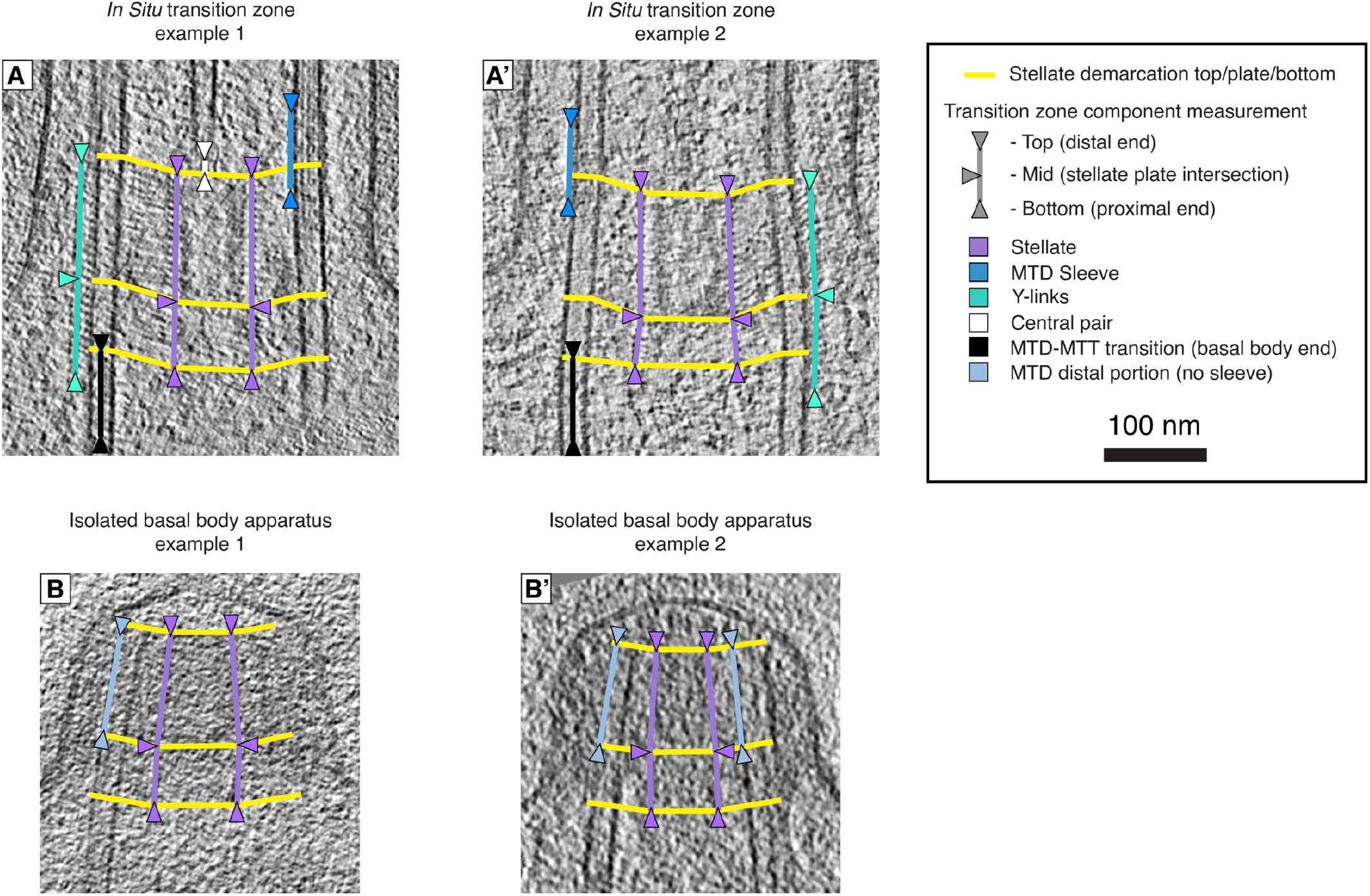
Methodology for manually measuring each TZ component in cells **(A, A’)** and in basal bodies isolated immediately after ciliary abscission **(B, B’)**. The position and extent of each structure was measured relative to one or more clear demarcations of the stellate apparatus: the proximal end, the central plate, and the distal end. In the isolated basal bodies, note how MTDs lack the helical sleeve decoration and terminate immediately at the distal end of the stellate. Measurement of the cleaved MTD ends (labeled “MTD distal portion”) was used to position the site of flagellar autotomy (SOFA, Fig. 1B). The overlap of the SOFA with the *in situ* position of the MTD helical sleeve, as well as the absence of the sleeve from basal bodies after ciliary abscission, suggests that the sleeve structure may be involved in MTD severing.

**Figure S4.**
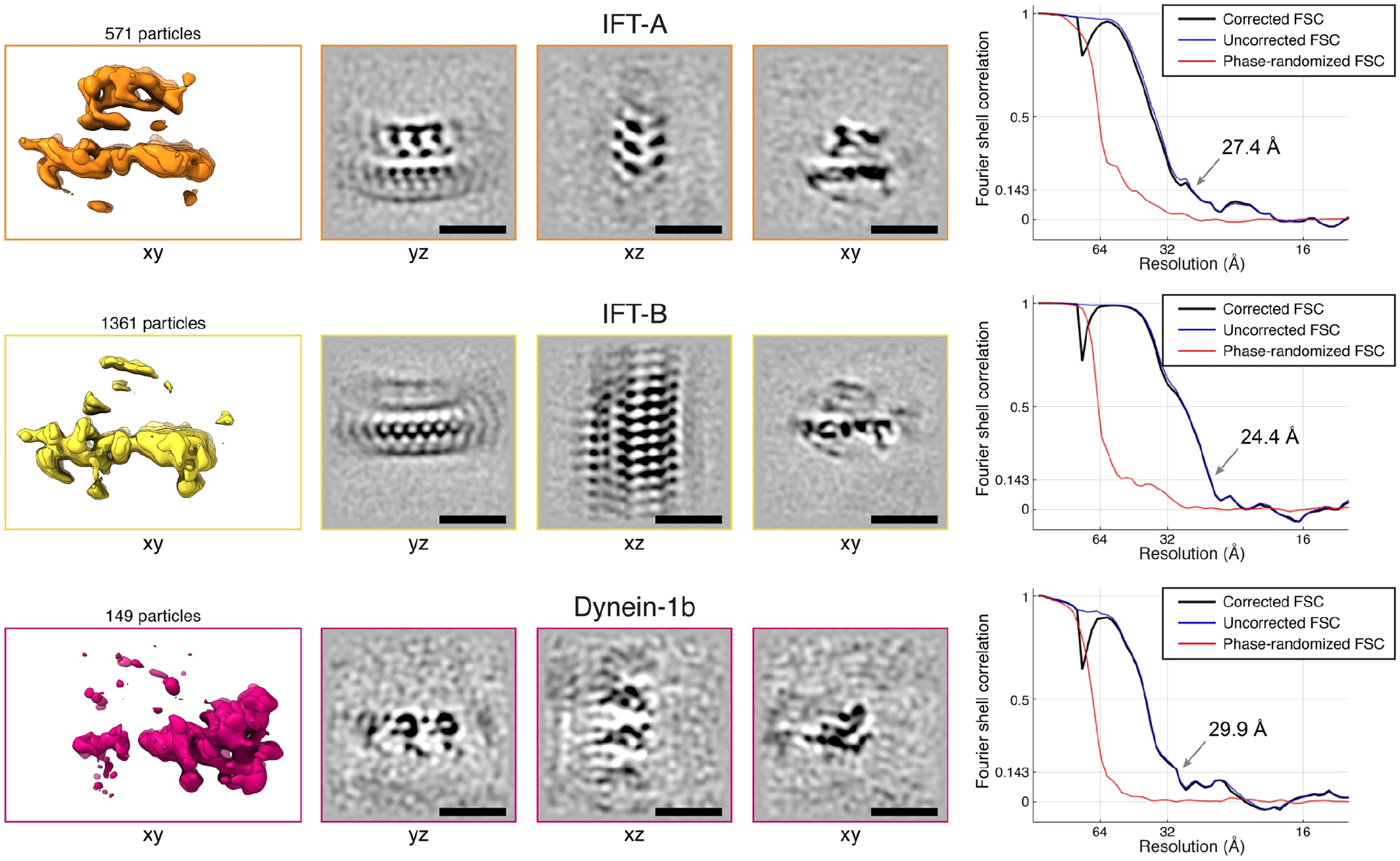
Individual subtomogram averages focused on different IFT train structures. Subtomogram averages focused on individual components of the IFT train: IFT-A (top row), IFT-B (middle row), and dynein-1b (bottom row). These averages were fused to generate the IFT model shown in Fig. 2. Left panels: 3D isosurface views, with the number of subvolumes (particles) in each average indicated. Central panels: 2D greyscale slices through the averages from three orientations (scale bars: 30 nm). Right panels: Fourier shell correlation curves showing the determined resolution of each average.

**Figure S5.**
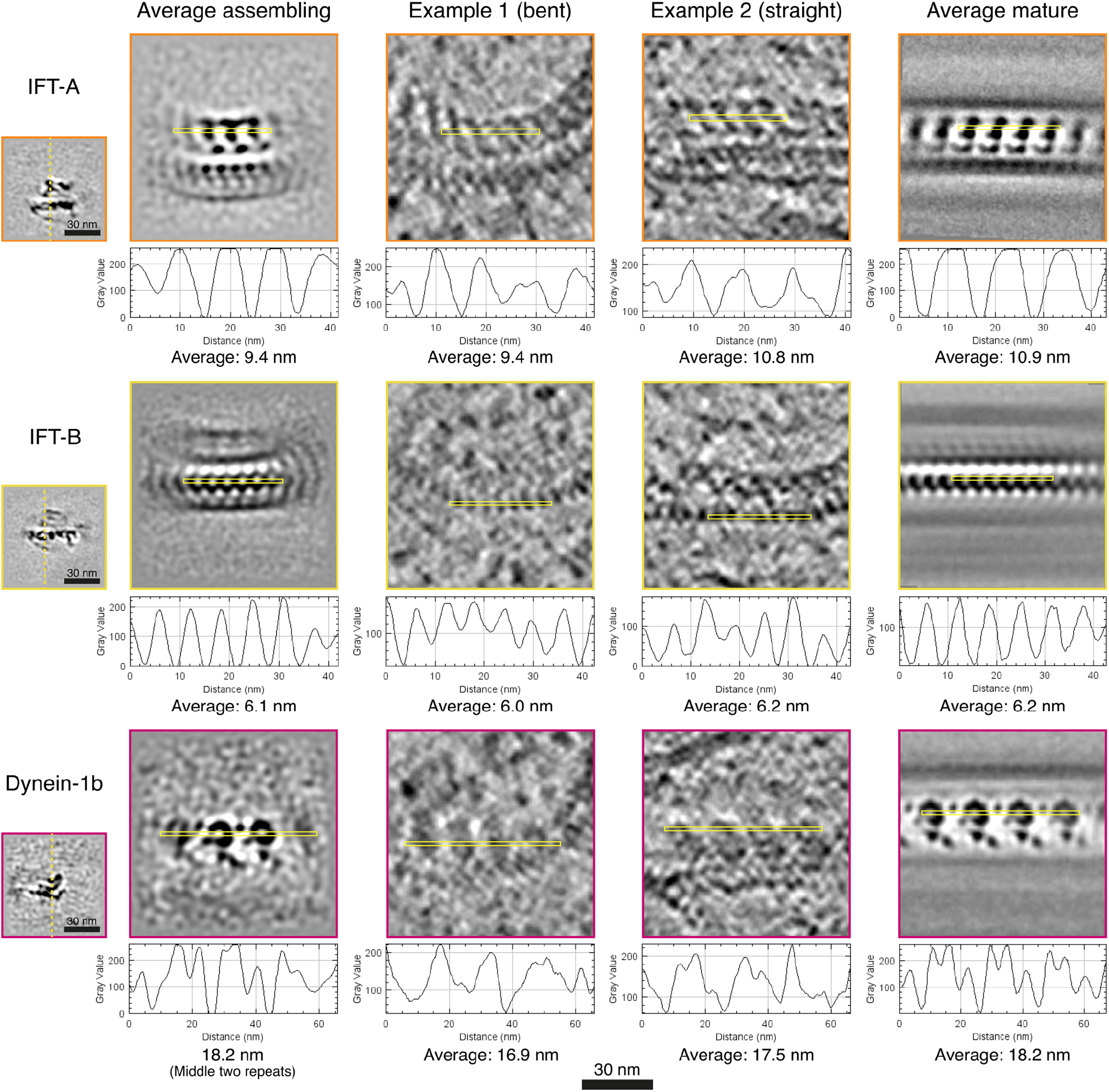
Periodicity of each IFT structure measured from greyscale 2D slices. For each structure, subtomogram averages of assembling trains are shown on the left, two examples of assembling trains from raw tomograms are shown in the middle, and subtomogram averages of mature trains (*14*) are shown on the right. The orientations of the longitudinal slices are indicated with dashed yellow lines in the cross-section views on the far left. Line scans along the yellow boxes in the longitudinal views were generated with FIJI software (*36*). Corresponding intensity plots are shown below each longitudinal slice, along with the average periodicity measured from each plot. Due to the flexibility and curvature of assembling trains in the cytoplasm, the IFT-A repeat is ~11 nm at its interface with IFT-B but only ~9.5 nm on the side that will face the membrane in mature trains after ciliary entry (see “average assembling” on left). In contrast, IFT-A in the cilium has an extended straight conformation with ~11-nm periodicity on both sides (see “average mature” on right). This difference between assembling and mature trains demonstrates the flexibility of IFT-A.

**Figure S6.**
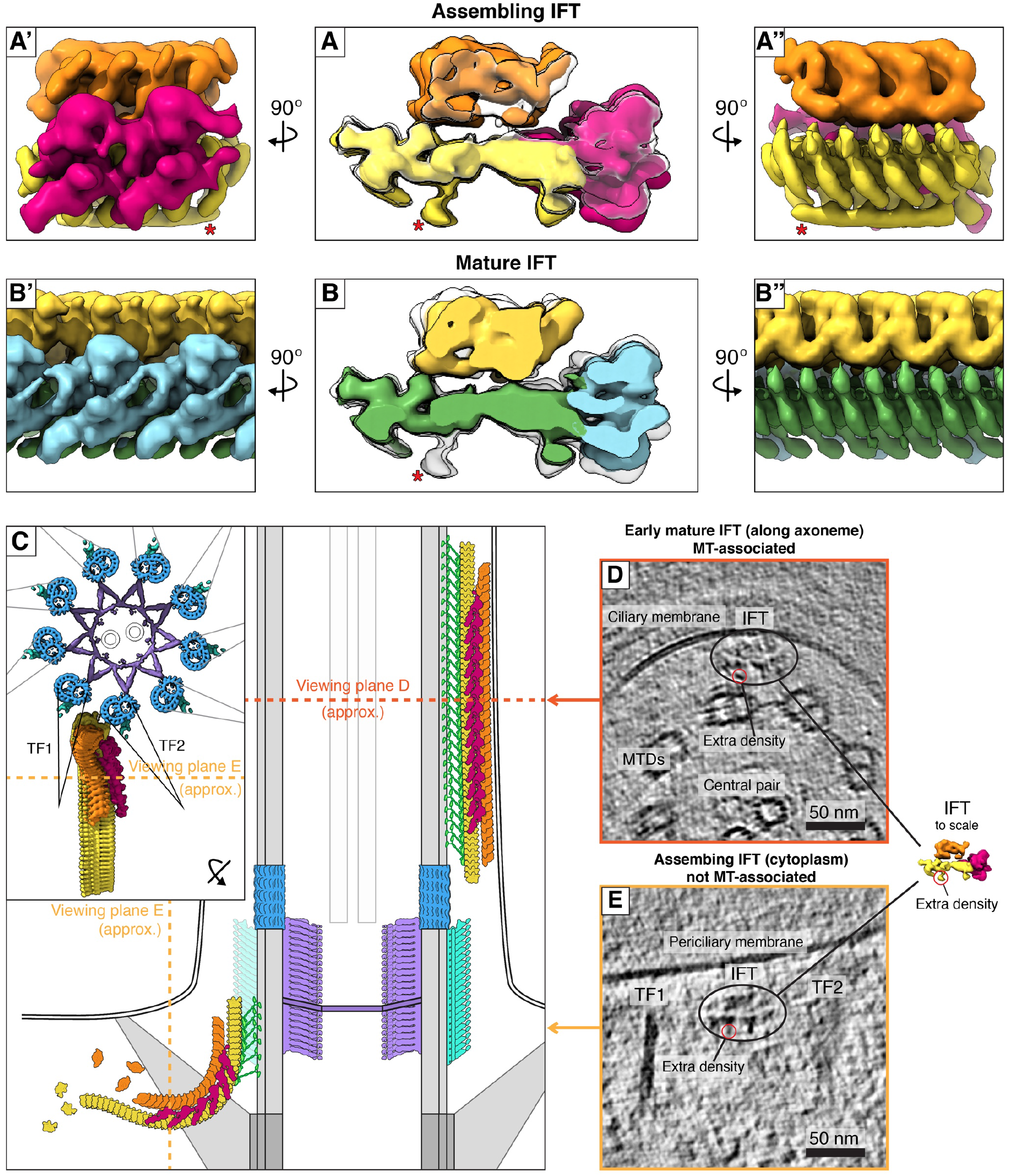
The extra IFT-B density on assembling trains. **(A)** 3D isosurface overlay between the subtomogram averages of the assembling IFT train at the ciliary base (colored as in Fig. 2) and the mature IFT train inside the cilium (from (*14*), white silhouette). Side views of the assembling train structure (without overlay) are shown in **A’** and **A’’**. The extra IFT-B density observed on assembling trains is indicated with a red asterisk. **(B)** 3D isosurface overlay between the structure of the mature IFT train (colored as in (*14, 37*)) and the assembling IFT train (white silhouette). Side views of the mature train structure (without overlay) are shown in **B’** and **B’’. (C-E)** The IFT-B extra density is clearly observed in 2D slices through raw tomograms (approximate viewing planes are diagrammed in **C**). **(D)** The extra density seen on a train that has just entered the cilium. Here, the extra density is positioned close to an unknown structure emanating from the MTD. **(E)** The extra density seen on a cytosolic assembling train passing between two transitional fibers (TF1 and TF2).

**Figure S7.**
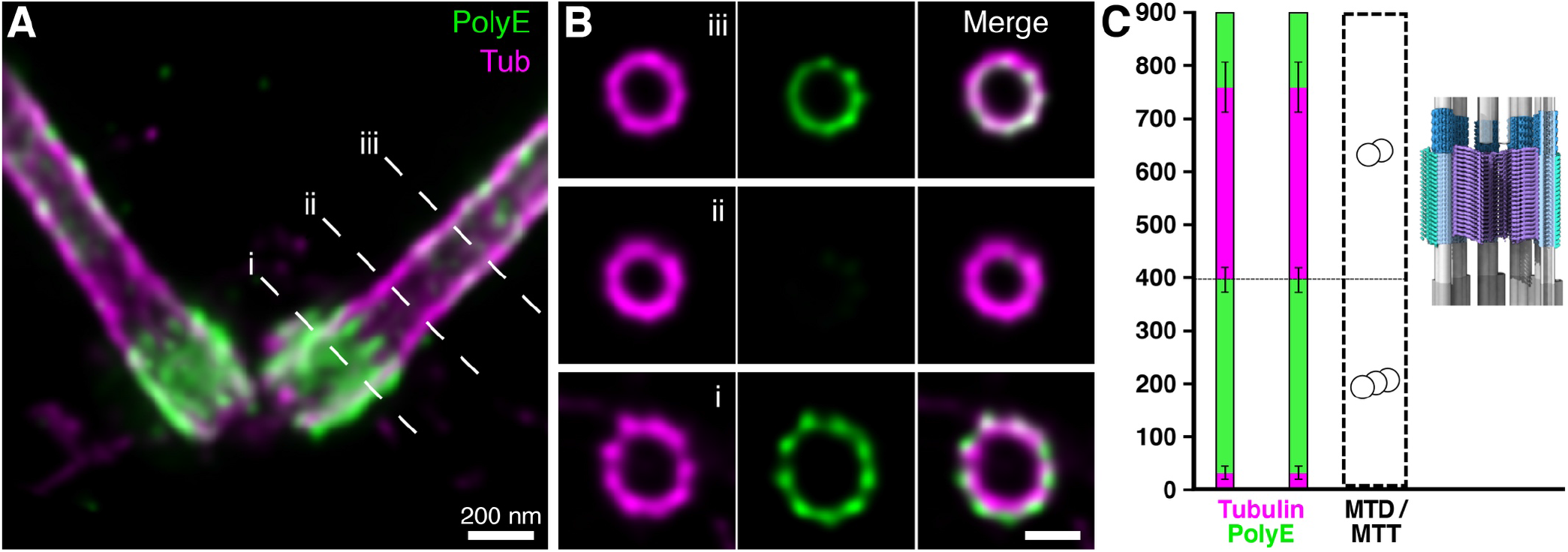
U-ExM shows a gap in PolyE staining at the TZ. **(A-C)** Tubulin staining is magenta, and PolyE staining is green. **(A)** Longitudinal overview and **(B)** corresponding cross-section views through the centriole (i), TZ (ii), and ciliary axoneme (iii). Signal for PolyE terminates at the distal end of the basal body, approximatively at the position of the MTT to MTD transition, leaving the TZ devoid of PolyE. PolyE signal reappears on the axoneme, distal of the TZ region. **(C)** Quantification of the PolyE signal from the proximal end of the centriole (0 nm on the Y-axis) through the TZ and the proximal ciliary axoneme. The cryo-ET model of the TZ is displayed to scale. Error bars show standard deviation. N= 30 centrioles and axonemes from 3 independent experiments. All scale bars: 200 nm.

**Figure S8.**
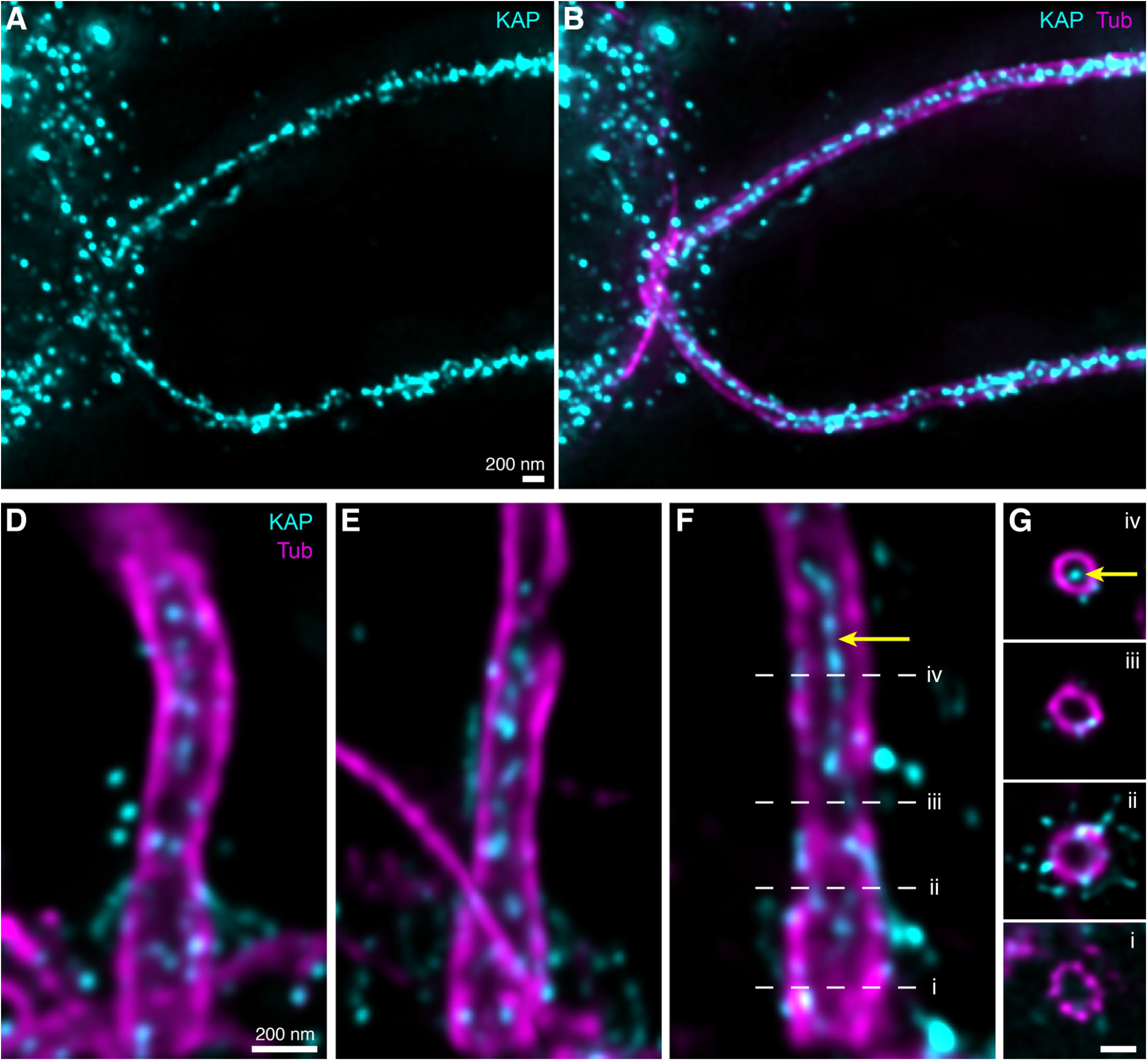
U-ExM reveals abundant KAP signal in the axoneme lumen, which is excluded from the TZ and centriole. Tubulin staining is magenta, and KAP-GFP staining is cyan. **(A-B)** Overview images of cilia extending from a *Chlamydomonas* cell. Panel A shows only the KAP-GFP signal, and panel B is a merge with tubulin. **(C-E)** Longitudinal views and **(F)** corresponding cross-section views through the centriole (i), proximal TZ (ii), distal TZ (iii), and ciliary axoneme (iv). The strong KAP-GFP signal in the axoneme lumen (yellow arrowhead) is excluded from the TZ and centriole. Presumably, the luminal KAP-GFP protein cannot diffuse past the stellate plate (Fig. 4H). All scale bars: 200 nm.

## Methods

### Cell culture

*C. reinhardtii* cells were obtained from the Chlamydomonas Resource Center (University of Minnesota, Minneapolis, MN USA). For *in situ* cryo-ET, we used strain CC-3994 (*mat3-4* mt+) (*38*). This strain has smaller cells, which is beneficial for vitrification during plunge freezing and also increases the chances of hitting a transition zone region by FIB milling. For cryo-ET of basal bodies following ciliary abscission, we used strain CC-4533 (*cw15* mt-; Jonikas CMJ030) (*39*). This strain lacks a cell wall and thus easily ruptures during blotting onto EM grids, enabling cryo-ET of basal bodies in thin ice without FIB milling. Cells were induced to form gametes by nitrogen starvation in order to reduce the size of the cell body. Ciliary abscission occurred spontaneously during blotting (no deciliation treatment was used), only seconds before cryo-immobilization by plunge freezing. For U-ExM, we used *C. reinhardtii* strains expressing IFT46-YFP (*40*), KAP-GFP (CC-5408) (*32, 41*), and D1bLIC-GFP (CC-4488) (*42*). For all experiments, cells were grown in Tris–acetate–phosphate (TAP) medium, with constant light, normal atmosphere, and room temperature.

### Cell vitrification and cryo-FIB milling

Both vitrification and FIB milling protocols were performed as described in (*17, 19*). Using a Vitrobot Mark 4 (FEI Thermo Fisher), 4 μL of cell culture (mid-log phase, diluted to ~1,000 cells per μL) was blotted onto holey carbon-coated 200-mesh copper EM grids (Quantifoil Micro Tools) and plunge-frozen using a liquid ethane/propane mixture. CC-3994 and CC-4533 cells were blotted onto R2/1 and R3.5/1 grids, respectively. Cryo-FIB sample preparation was performed using either a Quanta or Scios dual-beam FIB/SEM instrument (FEI, Thermo Fisher Scientific). Mounted in an Autogrid support, the grids were first coated with an organometallic platinum layer using a gas injection system. Subsequently, the cells were milled with a gallium ion beam to produce ~70- to 200-nm-thick lamellas, exposing the cellular interior.

### Cryo-electron tomography

EM grids with FIB-thinned cells were transferred to a 300 kV Titan Krios microscope (FEI, Thermo Fisher Scientific), equipped with a post-column energy filter (Gatan) and a K2 Summit direct detector camera (Gatan). Using SerialEM software (*43*), tilt-series were acquired from −60° to +60°, with 2° increments (bidirectional, separated at −0° or −20°). A subset was collected using a dose-symmetrical tilt scheme (*44*). Images were recorded in movie mode at 12 frames per second, with an object pixel size of 3.42 Å (magnification of 42,000x) and a defocus of −4 to −6 μm. The total accumulated dose for each tilt-series was ~100 e−/Å^2^. Each tomogram was acquired from a separate cell and therefore can be considered a biological replicate. Several different cell cultures and >10 imaging sessions were used to produce the dataset of 19 tomograms.

### Tomogram reconstruction

Prior to reconstruction, frames from the K2 detector were drift-corrected with MotionCor2 software (*45*) using 3 × 3 patches, after which tilt-series stacks were assembled using the TOMOMAN pipeline (https://github.com/williamnwan/TOMOMAN). The defocus of each tilt was estimated using Gctf (*46*), tilt images were low-pass filtered proportional to the accumulated dose (“dose-filtering” in TOMOMAN), and then CTF-correction was applied using the CTFphaseflip tool in IMOD software (*47*). Using IMOD, the tilt-series were aligned with patch tracking and reconstructed with weighted back-projection to generate tomographic volumes. We applied quality control criteria including tilt-series alignment scores and the power spectra of individual tilts to omit poor tilts from tomograms. Contrast enhancement for display purposes (Figs. 1A, 2B-C, S6) and distance measurements (Figs. S2, S3, S5) was achieved using the tom_deconv deconvolution filter (https://github.com/dtegunov/tom_deconv) (*48*).

### Particle picking and subtomogram averaging

IFT-A, IFT-B, and dynein-1b positions on each IFT train, as well as the stellate fibers, Y-links, and MTD sleeve, were manually picked along the train or MTD in a consistent distal-to-proximal fashion, in deconvolution-filtered bin4 tomograms (13.68 Å pixel size) using IMOD’s 3dmod viewer. Subtomogram alignment and averaging was performed using STOPGAP (v0.3.1, https://github.com/williamnwan/STOPGAP/) (*49*). STOPGAP is a subtomogram averaging package that uses real-space refinement to iteratively align and average subtomograms. Initial positions for each type of particle were twice oversampled (compared to their roughly measured repeat lengths) and extracted along an interpolated line running through the filament’s corresponding particle positions in the CTF-corrected, dose-filtered tomograms. By aligning these particles to their own initial average, generated using randomized rotations along the filament’s longitudinal axis, we obtained bias-free *de novo* averages after particle positions and rotations converged. These averages were low-pass filtered to 40 Å and iteratively realigned using several cylindrical masks, tolerances and Euler angle sampling cones. Using the “place object” tool in UCSF Chimera software (*50, 51*), we visually inspected the relative orientations of particles at different cross correlation (CC) thresholds. We cleaned each average by omitting particles with aberrant orientations (and therefore low CC score) by setting a global CC score cutoff (Table S1). These low-scoring particles, as well as overlapping redundant particles from oversampling, were discarded using functions in the STOPGAP toolbox. The final round of alignment and averaging was performed with bin2 dose-filtered, CTF-corrected subvolumes (6.84 Å pixel size), using a wide cylindrical mask, a very tight cylindrical cross correlation mask (xyz translational sampling space), and a narrow Euler angle sampling cone. The resolution of the final averages (Table S1, Figs. S1, S4) was estimated using mask-corrected Fourier shell correlation (*52*) after splitting each set of particles into two half-sets (even and odd particle numbers) during the last alignment step.

**Table S1.** Parameters of the subtomogram averages. For each density map averaged in this study, this table lists the number of particles in the average, the subvolume box size, the global CC score cutoff, the resolution determined at FSC=0.5 and FSC=0.143 thresholds (Figs. S1, S4), and the repeat length of the structure (Figs. S2, S5).

| Particle type | Number of particles | Box size (px) | CC score cutoff (cleaning step) | Resolution @ FSC 0.500 (Å) | Resolution @ FSC 0.143 (Å) | Average repeat length (nm) |
| --- | --- | --- | --- | --- | --- | --- |
| IFT-A | 571 | 128 | 0.200 | 36.2 | 27.3 | 9.2 |
| IFT-B | 1361 | 128 | 0.100 | 28.6 | 24.4 | 6.2 |
| Dynein-1b | 146 | 128 | 0.350 | 38.2 | 29.9 | 18.1 |
| MTD Sleeve | 110 | 192 | 0.350 | 42.9 | 34.7 | 16.4 |
| Stellate | 206 | 256 | 0.140 | 34.6 | 28.2 | 8.2 |
| Y-link | 529 | 192 | 0.220 | 35.8 | 27.9 | 8.3 |

### Length measurements of transition zone components

Measurements of the transition zone components were performed using IMOD’s 3dmod viewer tool. Using the “slicer” window, we aligned each bin4 deconvolution-filtered tomogram so that the length of each desired structure was entirely contained within in the viewing plane. The length of a repetitive structure was recorded with the measuring tool, divided by the number of repeats, and multiplied by the pixel size (1.37 nm at bin4) to obtain the final repeat length. Each MTT/MTD/TZ structure contained within the tomographic volumes was measured separately, and these measurements were then averaged (Fig. 1B).

### Assignment of assembly state along mature and assembling IFT trains

After careful revision of the manually picked positions corresponding to IFT-A, IFT-B and dynein-1b along each IFT train, we used UCSF Chimera’s “place object” function to assign identities to each IFT-B subunit by noting if they were immediately adjacent to IFT-A (about one IFT-A per two IFT-B subunits) and/or dynein-1b (one dynein-1b per three IFT-B subunits). A caveat to the quantification of assembling trains in Fig. 3 could be a bias towards shorter trains, as the back ends of some longer trains were cut off by FIB milling (red “x” in Fig. 3A). IFT train location with respect to transition zone components was measured using IMOD’s 3dmod “slicer” tool. We made 2D slices through each bin4 deconvolution-filtered tomogram that displayed the front of each IFT train in plane with one of four easily recognizable TZ structures (top and bottom of the stellate apparatus, stellate plate, and the MTD-MTT transition point). The pixel distance between the transition zone component and the first IFT-B subunit of each train was noted and multiplied by pixel size (1.37 nm at bin4). This distance in nm could additionally be converted into number of IFT-B repeats, using the average IFT-B repeat length we observed (6.22 nm).

### Combination of averages

Individual averages of IFT components (Fig. S4) were fused to form a composite map of the complete assembling IFT train structure (Figs. 2D, S6A) using the UCSF Chimera “fit in map” tool. We used the full IFT-B average, which contained blurred densities of IFT-A and dynein-1b, as a framework upon which to fit segmented maps of the IFT-B, IFT-A and dynein-1b subcomplexes. We cross-checked the fit by also using the blurred densities of IFT-B in the full averages of IFT-A and dynein-1b to confirm that the parts were properly positioned with respect to each other. A similar approach was applied to assemble a composite map of the transition zone (Fig. 1C-E). Individual averages of the Y-links and stellate (Fig. S1) were fit together onto an MTD backbone (part of the stellate average) using the UCSF Chimera “fit in map” tool. Segmented single subunits of the Y-links, stellate, and MTD sleeve were then fit into the composite model and copied both longitudinally and helically, applying shifts and rotations (tom_rotate and tom_shift functions) using the Matlab-based TOM Toolbox (*53*). The number of repeats for each component was determined by measuring their repeat length (Fig. S2) and extent along the transition zones (Figs. 1B, S3).

### Tomogram visualization

Slices through tomographic volumes (Figs. 1A, 2B-C) were generated using the IMOD 3dmod viewer. Using the UCSF Chimera “Place Object” tool, 3D surface models of IFT and TZ structures were mapped back into the tomograms at the refined positions determined by subtomogram averaging (Figs. 2A-C). These models were then exported to ChimeraX software (*54*) for display. Subtomogram averages of individual complexes (Figs. 1F-H, 2D, S1, S4, S6) were also displayed in ChimeraX.

### Ultrastructure expansion microscopy (U-ExM) of *C. reinhardtii*

After growing the IFT46-YFP, KAP-GFP, and D1bLIC-GFP strains in TAP medium for three days, cells were sedimented on 12 mm coverslips coated with Poly-D-Lysine for 5 minutes. Following sedimentation, the U-ExM protocol was followed as previously described (*16*). Briefly, the 12 mm coverslips covered in unfixed *C. reinhardtii* cells were incubated in a solution containing acrylamide and formaldehyde in PBS for 5 hours at 37°C. TEMED (2.5 μl) and APS (2.5 μl) were added to monomer solution and vortexed before adding to the coverslips. The gelation proceeded for 5 min on ice, followed by 1 hr of incubation at 37°C. Samples embedded in the gel were then added to denaturation buffer (200mM SDS, 200mM NaCl, 50mM Tris-base) for 1.75 hr at 95°C. Denatured samples were then exposed to the first round of expansion overnight. The following day, gels were stained for 3 hr with anti-tubulin antibodies AA344 (β-tubulin, 1:400, scFv-S11B) (*55*) and AA345 (α-tubulin, 1:400, scFv-F2C) (*55*), in combination with either anti-GFP (1:200, Torrey Pines Biolabs, TP401) or anti-PolyE (1:500, AdipoGen, AG-25B-0030). After 30 min of washing in PBS with 0.1% Tween-20 (PBST), gels were incubated for 3 hr with a secondary antibody solution containing goat anti-rabbit Alexa Fluor 488 IgG (1:400, Thermo Fisher Scientific, A11008) and goat anti-mouse Alexa Fluor 568 IgG H +L (1:400, Thermo Fisher Scientific, A11004). All immunolabeling steps were carried out at 37°C with gentle shaking. Following secondary antibody incubation, gels were washed for 30 min in PBST, and expanded overnight in ddH_2_O. The final expansion factor varied from 4.2x to 4.6x, determined by using a caliper to measure the dimensions of the expanded gel.

### Fluorescence microscopy

Imaging was performed using either an inverted Leica TCS SP8 confocal microscope or a Leica Thunder DMi8 widefield microscope. Confocal Z-stacks were acquired with a 63x 1.4 numerical aperture oil objective (using Lightning dual-color deconvolution), a step size of 0.12 nm, and a pixel size of 35 nm. Widefield Z-stacks were acquired with a 63x 1.4 numerical aperture oil objective (using the Thunder Small Volume Computational Clearing mode), a step size of 0.14 nm, and a pixel size of 100 nm.

### Fluorescence microscopy data analysis

Image analysis and maximal intensity projections were performed using Fiji software (*36*). PolyE coverage (Fig. S7C) was measured by generating maximal projections of side-view basal bodies and measuring the full width-half maximum of the PolyE signal compared to the tubulin signal. For semi-automated IFT tracing, we used the Fiji Simple Neurite Tracer (SNT) plugin (*56*) with default settings and the “A* search” algorithm. Z-stacks of equivalent orientation and dimensions for IFT46-GFP, KAP-GFP, and D1bLIC-GFP were cropped to equal dimensions and individually opened in the SNT plugin. Using Z-stacks with obvious train-like structures, strongly stained points along the train were selected in 3D. The semi-automated software traced the complete train, and faithful tracing was validated by eye. The tracing profiles of each train were superimposed into one image (Fig. 4F). To convert train lengths from pixels to nanometers, we calculated local expansion factors using the basal body attached to each train as an internal ruler (basal body proximal end = 225 nm), as previously measured by *in situ* cryo-ET (*19*). Train lengths were then plotted and statistically analyzed using Graphpad Prism 8.

## Acknowledgments

We thank Dennis Diener and Karl Lechtreck for helpful discussions. G.P. and P.A.V. thank Tobias Furstenhaupt from the MPI-DBG EM facility for instrument support. This work was funded by the European Research Council (ERC) under the European Union’s Horizon 2020 research and innovation program (Grant 819826) to G.P., the Swiss National Science Foundation (SNSF) Grant PP00P3_187198 to P.G., and the ERC ACCENT Starting Grant 715289 to P.G. Additional funding for personnel and instrumentation was provided by the Helmholtz Zentrum München and the Max Planck Society.

## Author contributions

M.S., P.S.E., and B.D.E acquired cryo-ET data of FIB-milled *C. reinhardtii* cells. G.A.V. prepared samples and acquired cryo-ET data of isolated basal bodies. M.A.J. prepared samples and acquired cryo-ET data of intact cilia. H.v.d.H. analyzed all the cryo-ET data, with help from M.A.J. and W.W. U-ExM data acquisition and analysis was performed by N.K., with guidance from P.G. and V.H. Additional mentorship was provided by W.B. and J.M.P., along with access to instrumentation. G.P., V.H., P.G., and B.D.E. conceived and supervised the study. H.v.d.H., N.K., and B.D.E. wrote the paper, with input from all authors.

## Data and materials availability

Cellular tomograms, subtomogram averages, and composite maps will be deposited the Electron Microscopy Data Bank (EMD-####). All data needed to evaluate the conclusions of the study are present in the paper and the supplementary materials. Additional data is available from the authors upon request. Correspondence and requests for materials should be addressed to B.D.E.

